# Efficiency and precision of miRNA biogenesis modes in plants

**DOI:** 10.1101/388330

**Authors:** Belén Moro, Uciel Chorostecki, Siwaret Arikit, Irina P. Suarez, Claudia Höbartner, Rodolfo M. Rasia, Blake C. Meyers, Javier F. Palatnik

## Abstract

Many evolutionary conserved microRNAs (miRNAs) in plants regulate transcription factors with key functions in development. Hence, mutations in the core components of the miRNA biogenesis machinery cause strong growth defects. An essential aspect of miRNA biogenesis is the precise excision of the small RNA from its precursor. In plants, miRNA precursors are largely variable in size and shape and can be processed by different modes. Here, we optimized an approach to detect intermediates during miRNA biogenesis. We characterized a miRNA whose processing is triggered by a terminal branched loop. Plant miRNA processing can be initiated by internal bubbles, small terminal loops or branched loops followed by dsRNA segments of 15-17 bp. Interestingly, precision and efficiency vary with the processing modes. Despite the various potential structural determinants present in a single a miRNA precursor, DCL1 is mostly guided by a predominant structural region in each precursor in wild-type plants. However, our studies in *fiery1, hyl1* and *se* mutants revealed the existence of cleavage signatures consistent with the recognition of alternative processing determinants. The results provide a general view of the mechanisms underlying the specificity of miRNA biogenesis in plants.

## Introduction

MicroRNAs (miRNAs) are endogenous small RNAs found in plants and animals. In plants, miRNA primary transcripts are transcribed by RNA polymerase II, capped, spliced and polyadenylated [reviewed in (1–3)]. They are processed in the nuclei by complexes that cut the precursor using RNAse type III activity to release a miRNA/miRNA* duplex. The ~21 nt miRNA is methylated and loaded into an ARGONAUTE (AGO) protein, generally AGO1 (1, 2). MiRNAs provide AGO complexes with sequence specificity to identify target RNAs by base complementarity; the target RNAs may be cleaved, translationally repressed or destabilized (1, 2).

Transcription factors can control the expression of specific *MIRNAs,* while general factors such as Elongator complex have more global effects on miRNA levels (1). Resulting *MIRNA* primary transcripts are then recognized by a dicing complex, that harbors DICER-LIKE1 (DCL1) and the core components HYPONASTIC LEAVES 1 (HYL1), a dsRNA binding protein (4–7) and the zinc finger domain SERRATE (SE) (8–11). DCL1, HYL1 and SE interact *in vivo* and are required for an efficient and precise processing of miRNA precursors (12–17). Several other factors have been shown to aid on miRNA biogenesis modulating the final levels of the small RNAs [reviewed in (1)].

Plant *MIRNA* precursors comprise a largely variable group of stem-loop shapes and sizes (18, 19), contrasting with the stereotypic precursors found in animals (20). Interestingly, *MIRNA* processing can occur through different modes (21). In some cases, the miRNA processing machinery recognizes a 15-17 bp dsRNA stem below the miRNA/miRNA* duplex and above an internal bubble to produce a first cut that releases a stem-loop, resembling miRNA processing in animals (22–25). In other cases, a first cut by DCL1 is produced in the distal part of the precursor, below a small terminal loop, and precursor processing proceeds in a loop-to-base direction (19, 21, 26). In either case, after the first cleavage reaction, DCL1 produces a second cut ~21 nt away from the first cut releasing the miRNA/miRNA* duplex. DCL1 can also continue producing further sequential cuts every ~21 nt generating several small RNA duplexes (19,26–28). Intriguingly, precursors that harbor a terminal branched loop can be recognized by the DCL1 processing complex to generate unproductive cuts (29). Cleavage triggered by the recognition of terminal branched loops has been linked to a pathway leading to the destruction of the precursors (29).

We have previously described a method called SPARE (Specific Parallel Amplification of RNA ends) to identify miRNA processing intermediates (21) that was mainly used in *fiery1* mutants as they accumulate remnants of *MIRNA* processing due to a low XRN activity (30, 31). Here, we optimized the method and describe the processing of many previously uncharacterized *MIRNAs.* Almost one third of the evolutionarily conserved *MIRNAs* are processed from the loop to the base. Furthermore, we describe the biogenesis of *MIR157c,* which depends on the recognition of a terminal branched loop, a process previously shown to eliminate *MIRNAs*. We found differences in the efficiency and precision among the processing modes of Arabidopsis, which in can in turn the sequence of the small miRNAs. Although in wild-type plants *MIRNA* processing proceeds through the recognition of a main structural determinant in most cases, alternative structures may become relevant under certain conditions.

## Material & Methods

### Plant Material

*Arabidopsis thaliana* used in the study were wild-type plants, accession Col-0, and plants defective in *HYL1, SE* and *FIERY1.* For the SPARE and small RNA library construction wild-type and mutant plants *hyl1; se* and *fiery1* were grown as described before (32)(1)(33). After that, 40 ten-day old seedlings were harvested and frozen in liquid nitrogen for each library. Seeds of *hyl1-2* (SALK_064863), *se-1* (CS3257) and *fiery1* (SALK_020882) were obtained from the Arabidopsis Biological Resource Center (ABRC). For the analysis of primary transformations *MIR157a, MIR157b and MIR157c* wild-type and mutant versions were used, grown under short day (8h light/16h dark) at 22 ºC.

### SPARE library construction

Total RNA was obtained using TRIzol reagent (Invitrogen). Independent biological duplicates were made for each genotype with 40 μg of total RNA as starting material. After DNase treatment (DNase I, NEB M0303S), RNA fragments including processing intermediates were ligated to an RNA oligo adapter as described before (21). Ligated RNA was purified using the Dynabeads™ mRNA DIRECT™ Purification Kit (Thermo Fisher Scientific, 61011) and divided into eight tubes for cDNA synthesis. The cDNA reactions were carried out with specific oligos using SuperScript™ III RT (Invitrogen, 18080093) and 14 or 15 *MIRNAs* in each tube. To do this, specific primers containing a 5’-common tail; 5’-TGGAATTCTCGGGTGCCAAGG-3’ were used (see Supplementary Table 1 for a complete list). The resulting cDNAs were pooled together for a single PCR reaction using Phusion^®^ High-Fidelity DNA Polymerase (NEB, M0530S) using FW1 5’-GTTCAGAGTTCTACAGTCCGA-3’ and RV1 5’-TGGAATTCTCGGGTGCCAAGG-3’ common primers. The PCR program was 98 ºC for one minute, 20 cycles of 98 ºC for 30 seconds, 63 ºC for 30 seconds, 72 ºC for 50 seconds and 72 ºC for 10 minutes. The PCR products were purified from an agarose gel. A second PCR was done, using the same conditions for ten cycles and Illumina indexed primers (Illumina RS-200-0012 TruSeq^®^ Small RNA Sample Prep Kit). After gel purification the PCR product was sequenced on the Illumina HiSeq platform at the University of Delaware.

### Raw data process and normalization

Reads were first processed to remove the adapters using cutadapt v1.14 (34) and only reads ranging from 18 to 51 nt in length after adapter removal were kept. Adapter-free reads were mapped to the miRNA precursors using Bowtie v1.1.2 (35). Reads counts were normalized by the linear count scaling method to allow comparison of reads from different libraries. The normalized abundance was calculated according to the formula: Normalized abundance = raw/(precursor match)*120,000.

### Analysis of cleavage sites in wild-type and mutant plants

Precursors with at least ten reads flanking the miRNA/miRNA* and detected in both biological replicates in wild-type plants were used for detailed analysis *(MIR156a, MIR157c, MIR162a, MIR165a, MIR166a, MIR166b, MIR167a, MIR169a, MIR170, MIR171b, MIR171c, MIR393a, MIR398b, MIR398c, MIR399a, MIR399b, MIR399c, MIR408, MIR173, MIR400, MIR824* and *MIR825.* The same precursors were used for further analysis in *fiery1, hyl1* and *se* plants.

In base-to-loop precursors, the proximal cut is coincidental with the first cut, and the accurate position (most frequent cut) is defined as “0”. On the other hand, in loop-to-base precursors, the first cut (distal cut) and the second cut (proximal cut) are detected. Accurate positions were considered “0” in each case. Reads found in a range of +/− 3 nt from the “0” position were considered in our analysis. The shifts towards the miRNA were defined as −1, −2, and −3, respectively.

Primary transcripts were divided into three regions, the stem-loop, the 5’ and 3’ end, which are upstream and downstream regions, respectively. The stem-loop region was further classified in two regions, cuts flanking the miRNA/miRNA* duplex and other cuts. To calculate the frequency of cuts found in each region every read for each *MIRNA* was grouped into one of these categories.

### Small RNA libraries construction and analysis

1 μg of total RNA was extracted from wild-type seedlings using TRIzol (Invitrogen). The small RNA library was constructed using the Illumina TruSeq sRNA kit, and sequenced on the Illumina HiSeq platform at the University of Delaware. Only abundant miRNAs, with at least 100 reads of a small RNA in one biological replicate were considered for the isomiR frequency variation analysis.

### Bioinformatics Analysis

For the prediction of RNA secondary structure the software Mfold RNA folding (36) (http://unafold.rna.albany.edu/?q=mfold/RNA-Folding-Form) was used with its defaults parameters. Combined box/violin plots were made using the public software available on http://shiny.chemgrid.org/boxplotr/.

### *in vitro* transcription

Pri-miR157c and pri-miR171b were transcribed *in vitro* from plasmids containing a genomic copy of each *MIRNA* with T7 RNA polymerase. Before the transcription, primers containing the T7 RNA promoter on the 5’end and complementary to *MIR157c* and *MIR171b* were used for a PCR reaction. The expected PCR products were purified from a 2% (w/v) agarose gel. The transcription reaction was done in a two-step protocol. An initial mix of 10 μL template DNA (~900 ng/μL), 4 μL T7 RNA polymerase buffer and 6 μL DEPC treated water was heated at 95 °C for 2 minutes and then left for 15 minutes at room temperature. After that, a second mix was composed of the following: 20 μL 100 mM NTP each, 2.5 μL RNase Out (Promega), 3 μL T7 RNA polymerase (60 U, Thermo Fisher Scientific) and 38.5 μL DPEC treated water was added. The transcription reaction was carried out for 5 hours at 37 °C. The transcription products were purified from a denaturing 6% (w/v) polyacrylamide gel and precipitated (Tris 0.01 M pH=8, EDTA 0.001 M pH=8, NaCl 0.3 M and ethanol). 100 pmols of pri-miR157c and pri-miR171b transcript were 5’-end-labeled with ^32^P-γ-ATP (10 μCi/μL), using T4 polynucleotide kinase (Thermo Fisher Scientific). The reaction product were purified from a denaturing 8% (w/v) polyacrylamide gel using autoradiography and precipitated as described above.

### Nuclease digestions

Partial RNA digestions were performed with T1 (Fermentas, native and denaturing conditions), S1 nuclease (Fermentas), V1 RNAse (Ambion) and alkaline hydrolysis. The alkaline hydrolysis reaction was performed by heating at 95ºC for 3 minutes a mix of 1.5 μL RNA (1 μM) with 3 μL NaHCO_3_ (200 mM, pH=7). The reaction was stopped by adding loading buffer (80% v/v formamide, 0.025% p/v xylene cyanol, 0.025% p/v bromphenol blue, 20% v/v TBE and 50 mM EDTA pH=8) and freezing. The denaturing T1 digestion was done by heating 1.5 μL RNA (1 μM), 7.5 μL CEU buffer (20 mM sodium citrate pH=5, 1 mM EDTA and 7 M urea) at 50ºC for 5 minutes. After adding 1 μL of T1 (1 U/μL) the reaction went through 15 minutes at 50ºC and then was stopped by precipitating the RNA overnight. All the overnight RNA precipitations were performed by adding enough quantity of DEPC treated water for 100 μL, 10 μL NaCl (3 M), 300 μL ethanol and 1 μL of Glycoblue (Invitrogen). For the native T1 reaction, 1.5 μL RNA (1 μM),1 μL folding buffer (20 mM Tris pH=7.5, 25 mM KCl and 10 mM MgCl2), 0.3 μL of *E.coli* transfer RNA (Sigma, 0.1 mg/mL) and 6.2 μL DEPC treated water were incubated at room temperature for 15 minutes. After that, 1 μL T1 (0.01 U/μL) was added, the reaction was carried out for 15 minutes, and then stop by precipitating the RNA overnight. The S1 reaction was done by incubating for 15 minutes at room temperature a mix of 1.5 μL RNA (1 μM), 1 μL folding buffer, 0.3 μL *E.coli* transfer RNA (0.1 mg/mL) and 5.2 μL DEPC treated water. After that, 1 μL of S1 (1 U/μL) and 1 μL of ZnCl2 (10 mM) were added for a 15 minutes reaction and then stopped by precipitation of the RNA overnight. A second S1 reaction was performed as described before but with a higher enzyme concentration; 10 U/μL. For the V1 reaction 1.5 μL RNA (1 μM), 1 μL folding buffer, 0.3 μL *E. coli* transfer RNA (0.1 mg/mL) and 6.2 μL DEPC treated water were incubated for 15 minutes at room temperature, after which 1 μL V1 (0.001 U/μL) was added. The reaction was carried out for 15 minutes, and then stopped by precipitating the RNA overnight. A second V1 reaction was performed as described before but with a higher enzyme concentration; 0.01 U/μL. Finally the incubation control consisted of mixing 1.5 μL RNA (1 μM) and 8.5 μL DEPC treated water for 30 minutes at room temperature before precipitating overnight. To analyze the digested products, half of each sample was loaded on an 8% (w/v) denaturing polyacrylamide gel and run for 3 hours at 3000 V. The remaining half was loaded on a 6% (w/v) denaturing polyacrylamide gel and run for 4 hours at 3000 V. For pri-miR171b, only an 8% (w/v) denaturing gel was run, as described before. The results were visualized by Phosphor Imaging (Typhoon FLA 7000, GE).

### Transgenes

*MIR157a, MIR157b* and *MIR157c* were cloned from Col-0 genomic DNA. For the mutant versions of *MIR157c, MIR157c-ΔLS* and *MIR157c-SL* site directed mutagenesis was done as described before (21). Both the wild-type and mutant precursors were expressed from the *35S* promoter present in the CHF3 binary vector (37). Supplementary Table 2 lists the sequences of each vector.

### RNA expression analysis

12 μg of total RNA extracted from inflorescences using TRIzol (Invitrogen) was resolved on 17% (w/v) polyacrylamide denaturing gels (7 M urea). Each sample contained at least 20 independent primary transformants. For miR157 detection, an antisense primer was used. The probe was 5’end labeled with [γ-^32^P] ATP and hybridized as described previously (33). For the original image of the miR157 small RNA blot and auto-radiography see Supplementary Figure 1.

## Results

### Genome-wide identification of miRNA processing intermediates in wild-type seedlings

We modified the SPARE approach to efficiently detect miRNA processing intermediates in wild-type samples (Figure 1a, see Material & Methods and Supplementary Figure 2). First, we analyzed four samples from wild-type and *fiery1* of 10-day-old seedlings resulting in ~240,000 reads that mapped into miRNA precursors for each genotype (Appendix and Supplementary Table 3). Due to the relative position of the precursors’ specific oligos, the SPARE method detects the proximal cleavage site in precursors processed in a base-to-loop direction, and all the cleavage sites in those processed from the loop to the base (see Supplementary Figure 2 and Appendix). New data was obtained for 17 *MIRNAs,* including members of the *MIR399* family and *MIR173* that are processed in a base-to-loop direction (Figure 1b-e, and Appendix). These precursors harbor a structured dsRNA segment of 15-17 bp below the miRNA/miRNA* duplex (Figure 1b-e, and Appendix). As we analyzed seedlings, we did not detect precursors of the evolutionary conserved *MIR172* family, which are induced during Arabidopsis flowering (38–40).

**Figure 1.**
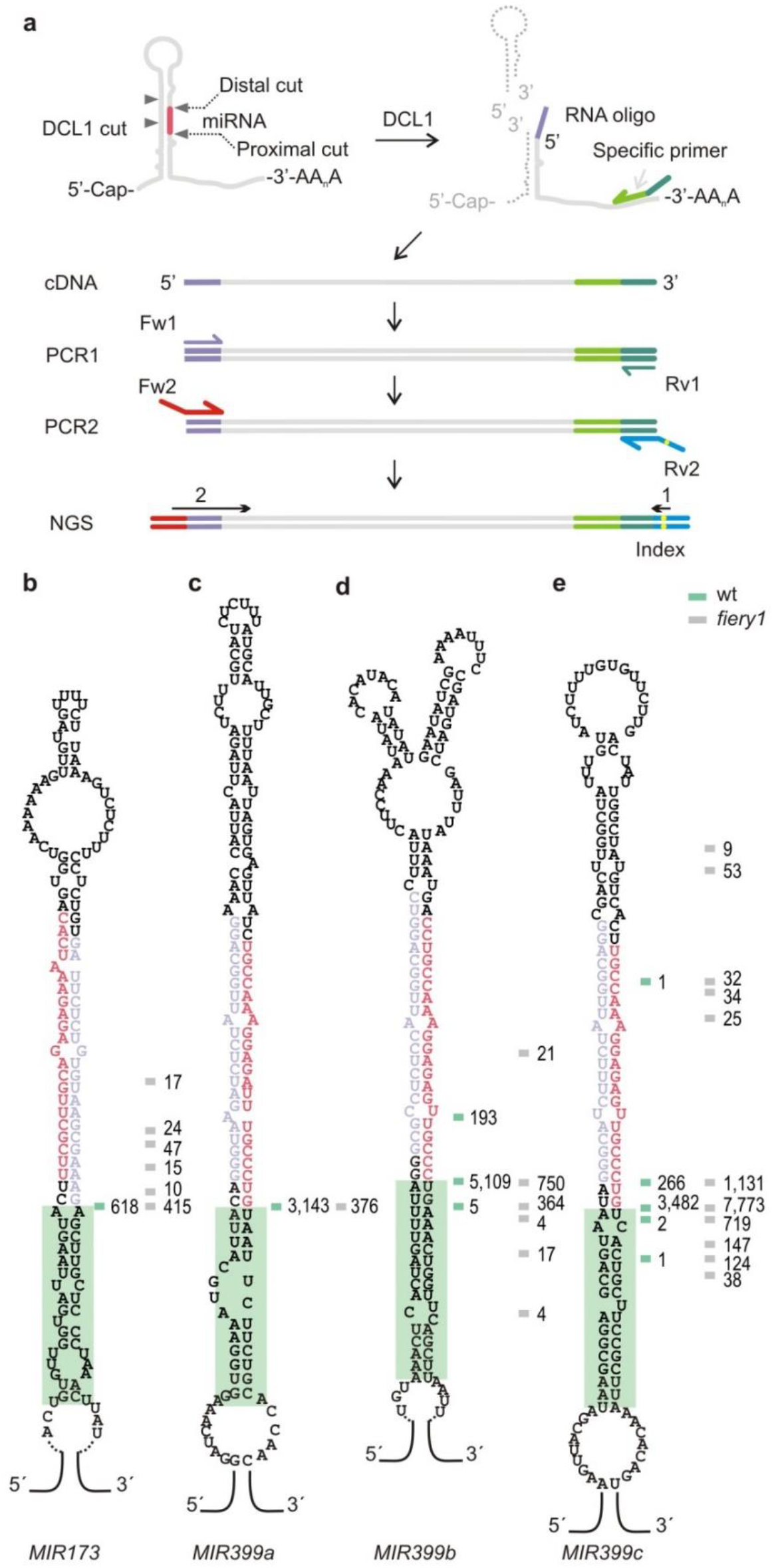
Detection of miRNA processing intermediates in wild-type plants. **(a)** Scheme of the SPARE method (See Supplementary Figure 2 for details). *MIRNAs* are cut by DCL1 (grey triangles) and the resulting fragments with free 5’ end are ligated to an RNA oligo (purple line) and subjected to retro-transcription (green arrow). DNA is sequenced after PCR amplification. **(b-e)** Predicted secondary structure of **(b)** *MIR173,* **(c)** *MIR399a,* **(d)** *MIR399b,* and **(e)** *MIR399c.* The miRNA is indicated in red and the miRNA* in light purple. Horizontal lines indicate cleavage sites detected in wild-type plants (green) and *fiery1* mutants (gray). Independent reads for each cut are shown as numbers next to the lines. A green box highlights a 15-17 bp dsRNA segment below the miRNA/miARN* duplex.

A caveat of small RNA sequencing data is that many mature miRNAs cannot be assigned to a unique precursor as they might originate from different *MIRNAs* of the same family. In our analysis of the SPARE data, we only considered reads longer than 30 nt, so that all sequences can be mapped unambiguously to specific precursors. One example is the case of *MIR399b* and *MIR399c,* which can potentially generate the same mature miRNAs (miRbase release 21). However, our analysis revealed that *MIR399b* and *MIR399c* precursors are specifically cleaved at different positions; generating two different miRNAs that are offset two nucleotides (Figure 1d-e).

### Identification of processing intermediates for precursors processed from the loop

We identified processing intermediates for additional *MIRNAs* processed from the loop, such as *MIR162a, MIR408* (Figure 2c-d). These precursors had two cuts flanking the miRNA/miRNA* (Figure 2a-d, and Supplementary Figure 2). A general analysis of the precursors processed by two cuts in a loop-to-base direction showed that the reads of the proximal and distal cut corresponded to approximately 50%, respectively, in wild-type plants. In contrast, 90% of the reads of in *fiery1* mutants corresponded to the position of the second cleavage site (Figure 2e, and Appendix). This result is consistent with the low XRN activity in *fiery1* that causes the accumulation of the remnants of miRNA biogenesis after the proximal cut (30, 31). Therefore, while *fiery1* increases the detection of certain RNA fragments, their relative accumulation might not reflect wild-type conditions. An example is *MIR408,* whose precursor has 99% of the reads in the second cut region in *fiery1* mutants, while in wild-type samples there are two cleavage sites yielding 20 and 80% of the reads (Figure 2d).

**Figure 2.**
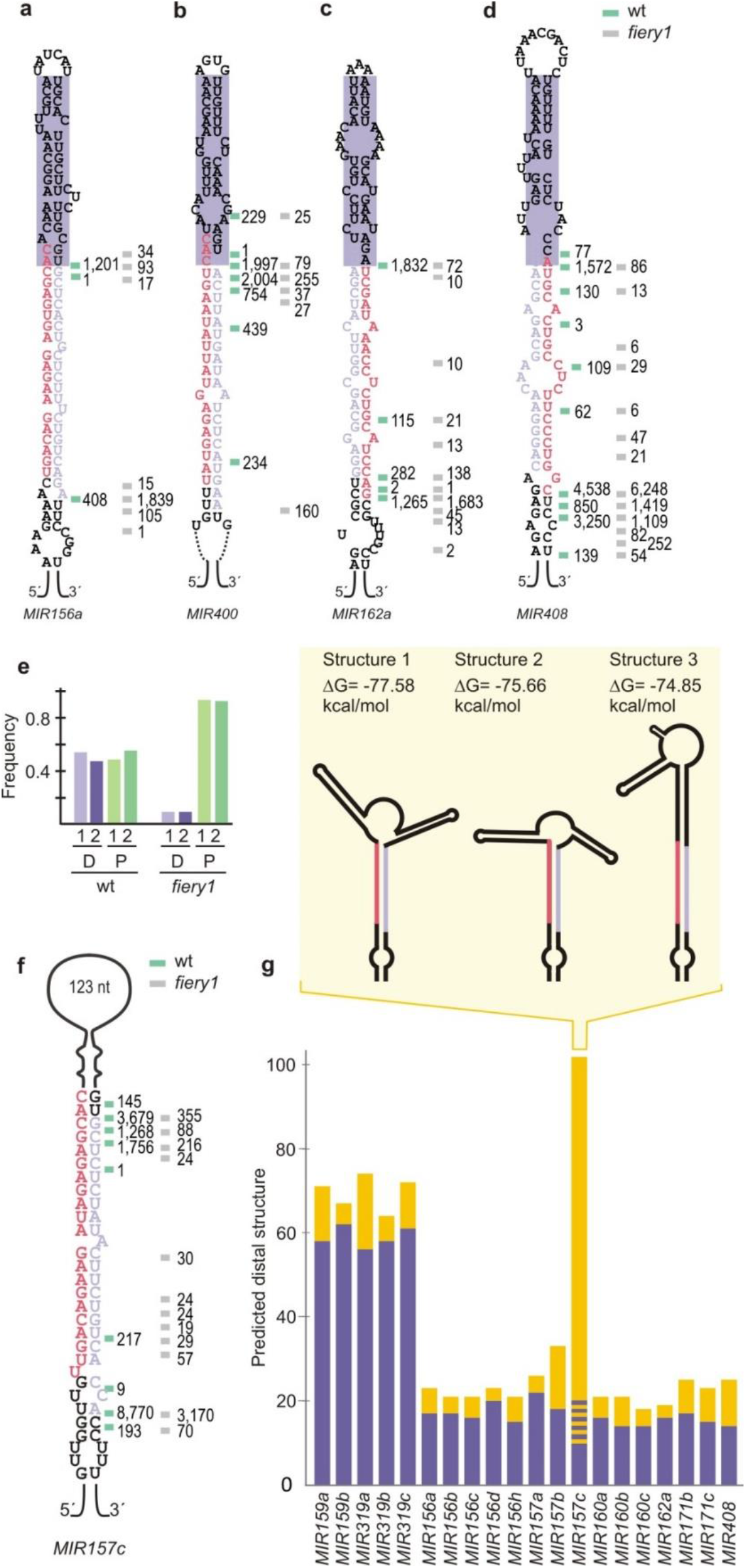
Processing intermediates of *MIR157c* and other *MIRNAs* processed from the loop. **(a-d)** Predicted secondary structure and processing intermediates of **(a)** *MIR156a,* **(b)** *MIR400,* **(c)** *MIR162a,* **(d)** *MIR408.* A purple box highlights a structured 15-17 bp dsRNA region above the miRNA/miRNA*. **(e)** Frequency of reads corresponding to the proximal (P) and distal (D) cuts in loop-to-base precursors in wild-type and *fiery1* mutants (the reads corresponding to two biological replicates are shown separately). **(f)** Processing intermediates of *MIR157c.* **(g)** Contribution of the terminal loop (in nucleotides, yellow bars) and the dsRNA segment above the miRNA/miRNA* duplex (in base pairs, purple bars) to the predicted distal structure of the precursors. A dashed line in *MIR157c* shows the variability of the secondary structure predictions for this precursor (see inset for a scheme and free energy of potential secondary structures).

Precursors processed by two cuts in a loop-to-base direction yield a 15-17 bp dsRNA segment above the miRNA/miRNA* followed by small terminal loops (Figure 2a-d, 2g) (21). An exception was the *MIR157c* precursor that has a large distal region of ~125 nt (Figure 2f-g). Unlike *MIR319/MIR159,* which are processed by several cuts and have a long dsRNA distal region (Figure 2g, and Appendix) (19, 21, 26), the distal part of *MIR157c* is mainly unstructured and it is processed by two cuts (Figure 2f and Supplementary Figure 3), suggesting that *MIR157c* has a particular biogenesis mode. RNA folding prediction showed several potential structures of similar energy, with different lengths of the dsRNA region above miR157/miR157* (inset Figure 2g, and Supplementary Figure 3).

### Productive recognition of a plant *MIRNA* via terminal branched loop

RNA folding prediction showed several potential structures of similar energy, with different lengths of the dsRNA region above miR157/miR157* (inset Figure 2g, and Supplementary Figure 3). To experimentally determine the secondary structure of the distal region of *MIR157c,* the precursor was transcribed *in vitro,* labeled on the 5’ end with ^32^P, treated with RNases that differentiate the structure of the RNA, and the products were resolved using in denaturing gels (Figure 3ab) (see Material & Methods for details). We found that S1 Nuclease, which selectively digests regions of ssRNA (41), could differentiate between the different predicted structures (Figure 2g, Figure 3a-c, and Supplementary Figure 3). The results show that *MIR157c* has a 17-19 bp dsRNA region above the miR157/miR157* followed by a terminal branched loop. Interestingly, the experimentally validated structure was not the most stable structure as predicted by the folding prediction algorithm. Previous results have shown that terminal branched loops can guide unproductive cuts that dampened the biogenesis of miRNAs (29). Instead, our results show that terminal branched loops can also trigger the productive biogenesis of small RNAs *in vivo.* In addition, we experimentally determined the secondary structure of the terminal region of *MIR171b* (Figure 3d-f). In this case, we found that the bases present in the small loop of *MIR171b* were detected as ssRNA (Figure 3d-e), correlating well with the secondary structure prediction.

**Figure 3.**
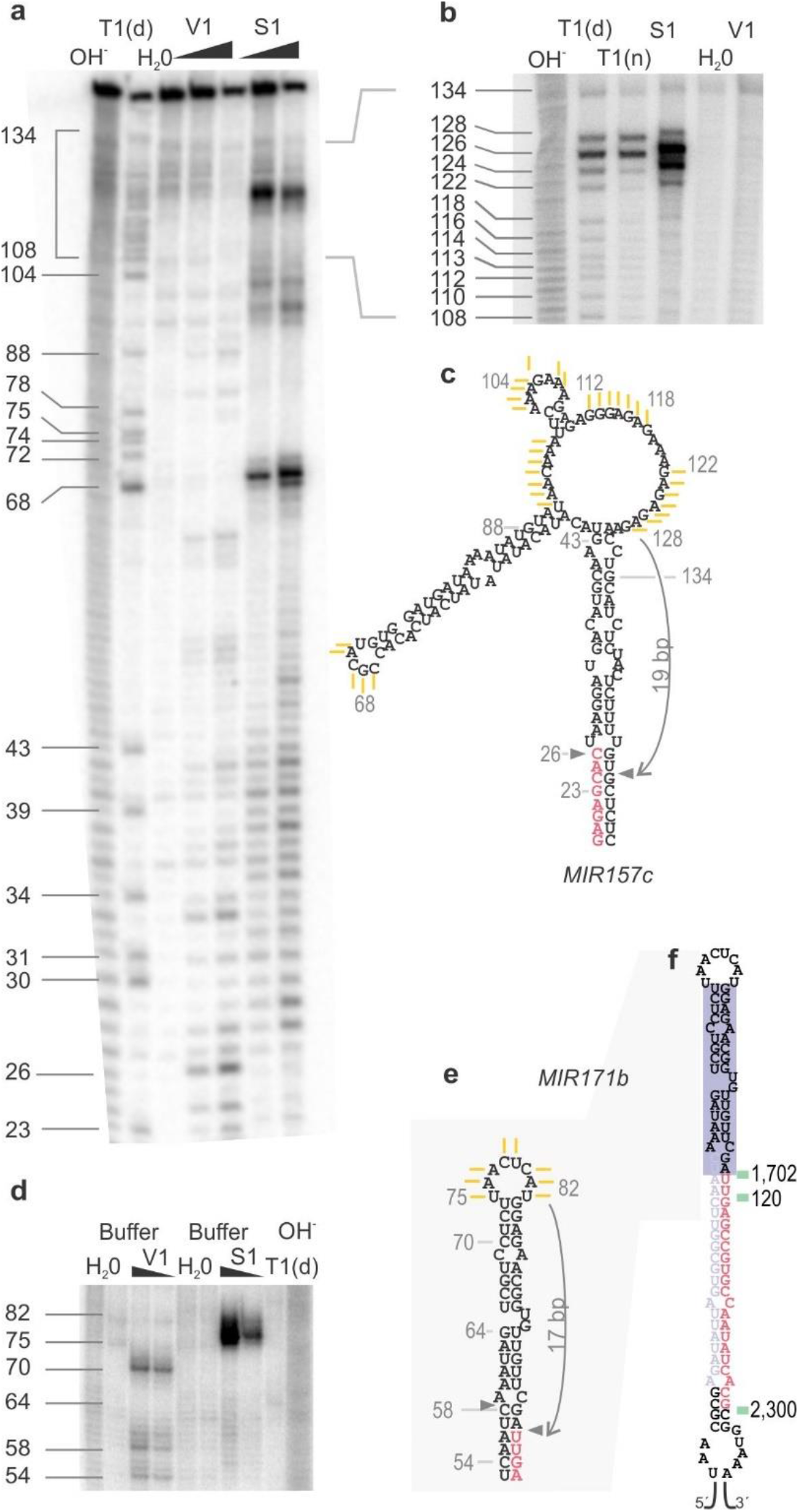
*MIR157c* is processed via a terminal branched loop. **(a-b)** Denaturing 8% **(a)** and 6% **(b)** (w/v) polyacrylamide gel of digested pri-miR157c. **(c)** Experimentally validated secondary structure of *MIR157c* distal region. Yellow lines represent the single stranded bases obtained after S1 digestion as determined from polyacrylamide gels **(a-b). (d)** Denaturing 8% (w/v) polyacrylamide gel of digested pri-miR171b. **(e)** Experimentally determined secondary structure of *MIR171b* distal region. Yellow lines indicate the single stranded bases obtained after S1 digestion as determined from polyacrylamide gels **(d). (f)** *MIR171b* processing patter obtained from wild-type plants (green lines). Notations are as indicated (a-b, d): OH-, alkaline hydrolysis, T1(d), T1 RNase in denaturing conditions and T1(n), T1 RNase in native conditions. The lines and numbers next to the gel correspond to bases on transcribed pri-miR157c and pri-miR171b, deduced from the T1(d) results. Two different V1 and S1 concentrations were used for each 8% (w/v) gel, while only the lowest concentration was employed on the 6% (w/v) gel. H_2_O, negative control. See Material & Methods for details.

### The terminal branched loop reduces the efficiency of miR157c biogenesis

To characterize the processing of *MIR157c,* we expressed different versions of the precursor from the *35S* promoter in Arabidopsis. First, we deleted the region below the miR157/miR157* duplex. We observed that the mutant precursor (MIR157c-ΔLS) accumulated similar levels of the miRNA with respect to the wild-type precursor (Figure 4a), demonstrating that these regions were dispensable for the biogenesis of miR157c. Next, we expressed the fold-back precursors of *MIR157a, MIR157b* and *MIR157c* from the *35S* promoter. All these *MIRNAs* are processed from the distal part of the precursor but differ in their terminal regions (Figure 2f and, Appendix). We found that the biogenesis of *MIR157a* and *MIR157b*, which have smaller loops, was more efficient than that of *MIR157c* (Figure 4b). Then, we replaced the terminal branched loop of *MIR157c* by a small loop of 4 nt (Figure 4c, *MIR157c-SL).* We found that *MIR157c-SL* was processed much more efficiently than the wild-type precursor. We further determined the phenotypic changes caused by the different *MIR157c* precursors on leaf number upon flowering (Figure 4d). In all cases, we found a good correlation between small RNA levels and the resulting phenotypes. Furthermore, the overexpression of the small loop version of *MIR157c* caused a significant increase in leaf number compared to the wild-type branched loop version (Figure 4d). Therefore, while precursors with branched loops can be processed productively to render mature miRNAs in plants, their processing efficiency seems to be relatively low.

**Figure 4.**
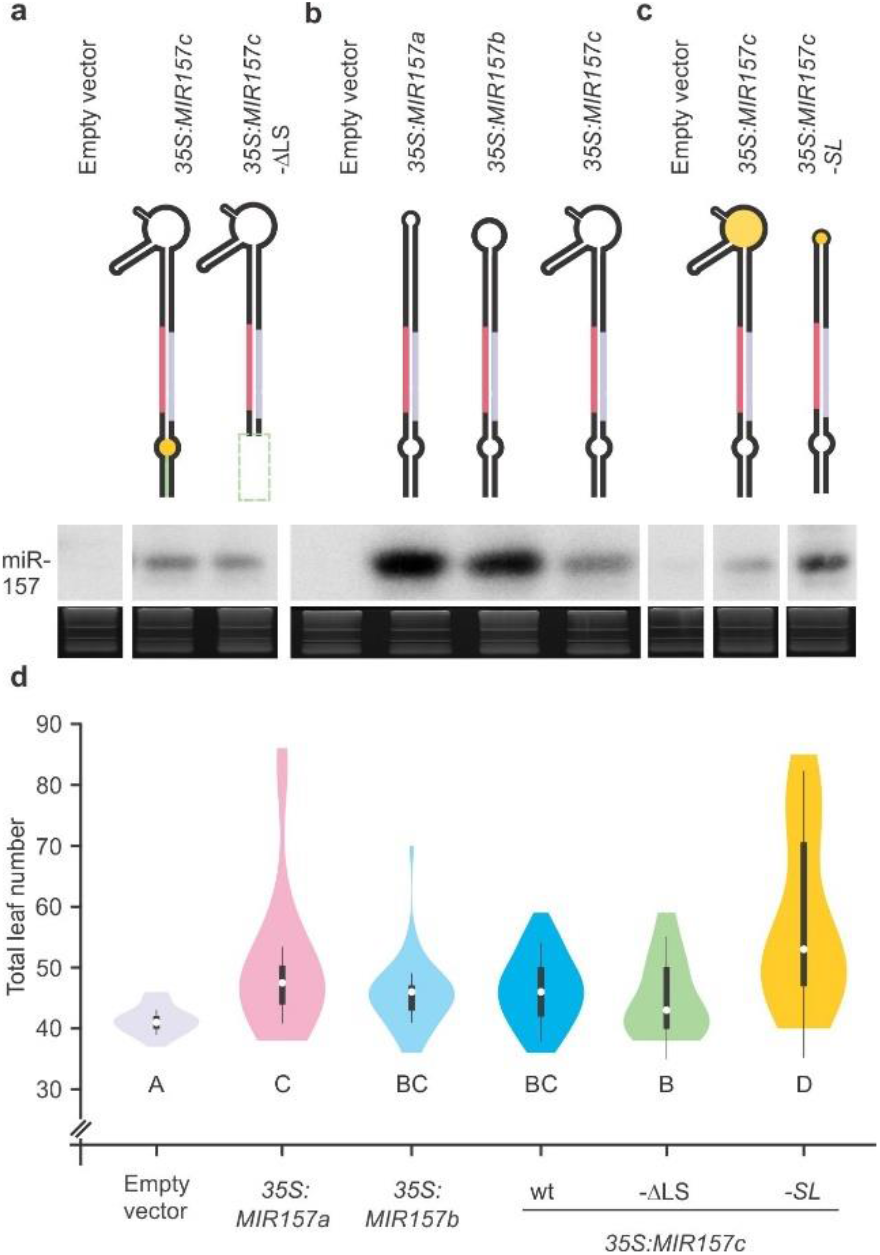
Modulation of *MIR157c* processing efficiency. **(a-c)** Small RNA gel blots of transgenic lines expressing different precursors from the *35S* promoter. Each sample represents a pool of 25 independent transgenic lines. Top panels show a schematic representation of the precursors analyzed. See Supplementary Table 2 for the expressed sequences of each vector. **(d)** Combined box/violin plots representing the distribution of rosette leaves number at the flowering time for primary transgenic plants over-expressing each construct grown in short days. Different letters indicate significant difference, as determined by Kruskal Wallis test, p<0,05. At least 50 primary transgenic plants were scored per construct.

### Flexibility in the processing of loop-to-base precursors generates variability in the miRNA sequences

While analyzing the processing intermediates, we realized that precursors processed from the base were accurately processed (Figure 1). In contrast, we noticed that there was flexibility in the position of the first cut in the precursors processed from the loop (Figure 2). To study the flexibility in the position of the first cut, we determined the most frequent cleavage position for the first cut on each precursor (position 0), and the three nearby positions on each side of this cut (positions +3, +2, +1, −1, −2, −3) (Figure 5ab). In wild-type plants, the precursors processed from the base have most of the cuts (>97%) in position 0 (Figure 5c), indicating an accurate processing of base-to-loop precursors. By comparison, we observed that ~84% of the cuts corresponded to position 0 in precursors that were processed from the loop (Figure 5d). Similar results were obtained in *fiery1* mutants, suggesting that the first cut in the precursors processed from the loop had more flexibility than in those processed from the base (Figure 5e-f).

**Figure 5.**
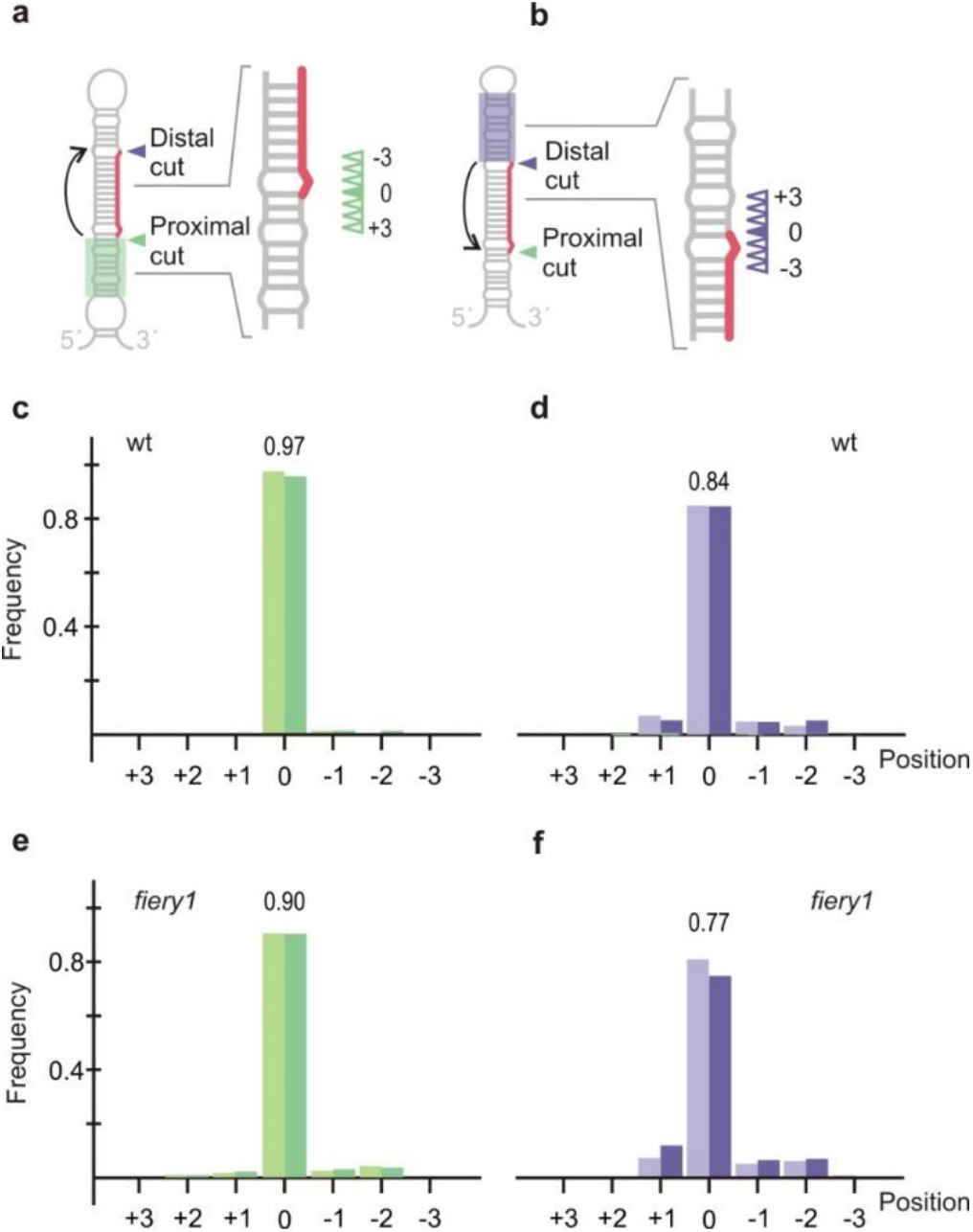
Accuracy of the first cut during *MIRNA* processing. **(a-b)** Schematic precursors processed in a loop-to-base **(a)** or base-to-loop **(b)** direction. The distal first cut is indicated with a purple triangle in **(a)**, while the proximal first cut is indicated with a green triangle in **(b)**. Position 0 is represented by a filled triangle, while the sloppy cuts (+3, +2, +1, −1, −2 and −3) are indicated with empty triangles. **(c-d)** Distribution of cuts for loop-to-base **(c)** and base-to-loop **(d)** precursors in wild-type plants. Data from two independent libraries are indicated by light and dark color bars. **(e-f)** Distribution of cuts for loop-to-base **(e)** and base-to-loop **(f)** precursors in *fiery1* mutants. Data from two independent libraries are indicated by light and dark color bars.

IsomiRs are miRNA variants that derive from the same precursor (42, 43). We investigated whether the flexibility in the processing of the precursors processed from the loop was translated into sequence heterogeneity of the mature miRNAs. To do this, we isolated and sequenced small RNAs from seedlings grown in similar conditions as those used for the SPARE analysis. Interestingly, we found a good correlation between the flexibility of the cuts detected in the precursors (Appendix, Figure 1–2) and the sequence variation of the small RNAs (Figure 6a-p, Supplementary Figure 4, and Supplementary Table 4). In general, miRNAs originated from precursors processed from the loop (Figure 6a-g) were more variable than those from base (Figure 6i-o). The average variation in the small RNAs due to flexibility on the position of the first cut turned out to be 14% in the miRNAs deriving from precursors processed from the loop (Figure 6h), while it was only 2% in those processed from the base (Figure 6p).

**Figure 6.**
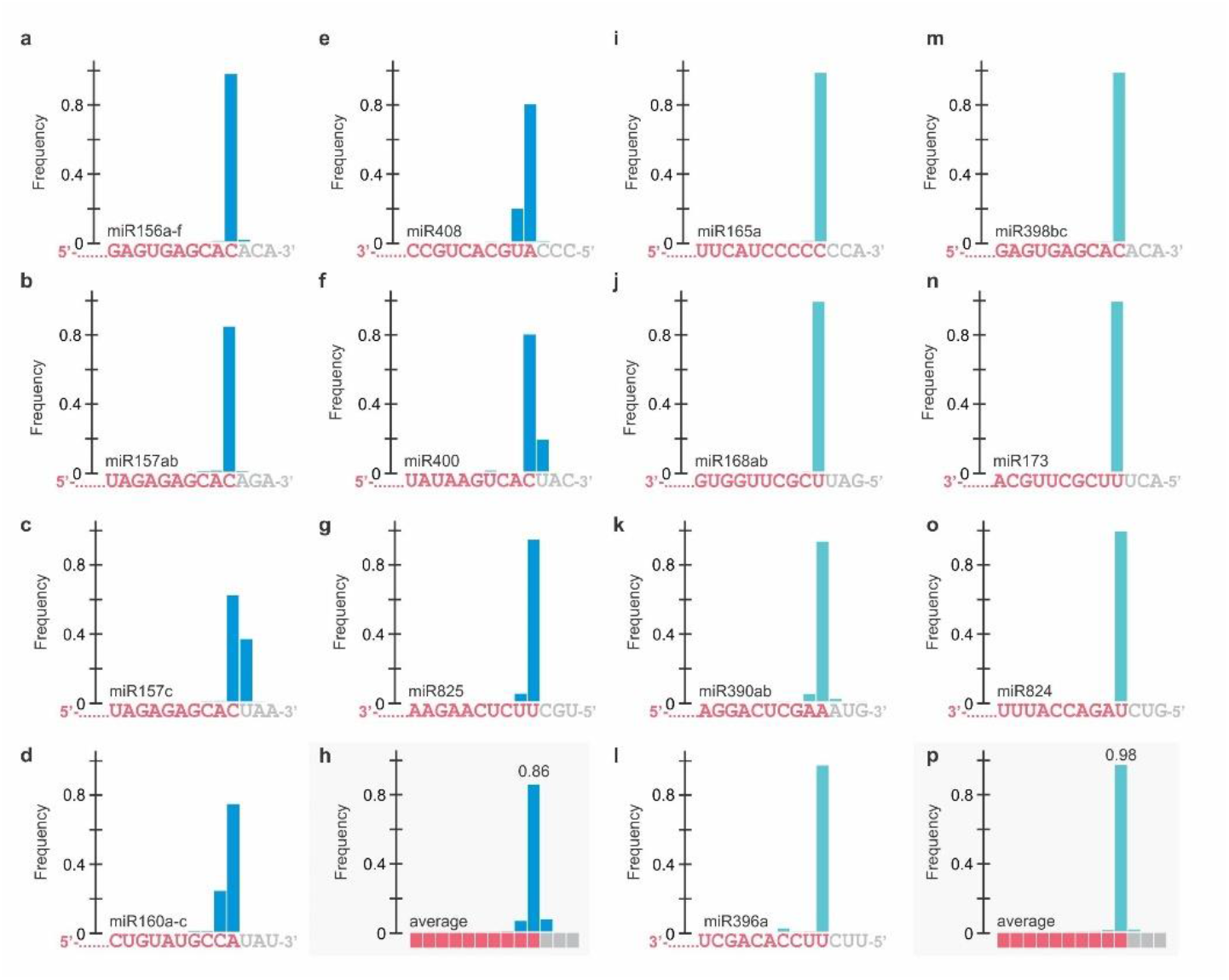
Small RNA sequence variation caused by variability in the position of the first cut. **(a-p)** miRNAs for which the biogenesis is loop-to-base are displayed on the left panels (blue bars) **(a-h)**, while base-to-loop miRNAs are displayed on the right panels (turquoise bars) **(i-p). (a)** miR156a-f, **(b)** miR157ab, **(c)** miR157c, **(d)** miR160a-c, **(e)** miR408, **(f)** miR400, **(g)** miR825, **(h)** average loop-to-base miRNA, **(i)** miR165a, **(j)** miR168ab, **(k)** miR390ab, **(l)** miR396a **(m)** miR398bc **(n)** miR173, **(o)** miR824, and **(p)** average base-to-loop miRNA. The miRNA sequence is indicated below each graph. The bar shows the frequency of the small RNAs ending at that position. For the average miRNA (h and p) the bases are depicted as squares.

### Precise processing of Arabidopsis precursors with diverse structures in wild-type plants

The genome-wide analysis performed on *fiery1* and wild-type plants [data obtained here, (21)] allows the determination of the processing direction of most Arabidopsis *MIRNAs* (Figure 7a). The optimization of the method performed here, allowed the efficient determination of precursors’ intermediates in wild-type plants, which do not have an enrichment of proximal cuts as seen in *fiery1* (Figure 2e). Interestingly, approximately 30% of the precursors are processed from the loop (Figure 7a). *MIRNA* processing can be triggered by small (Figure 7b) and branched (Figure 7c) terminal loops or internal bubbles (Figure 7d) followed by dsRNA regions.

**Figure 7.**
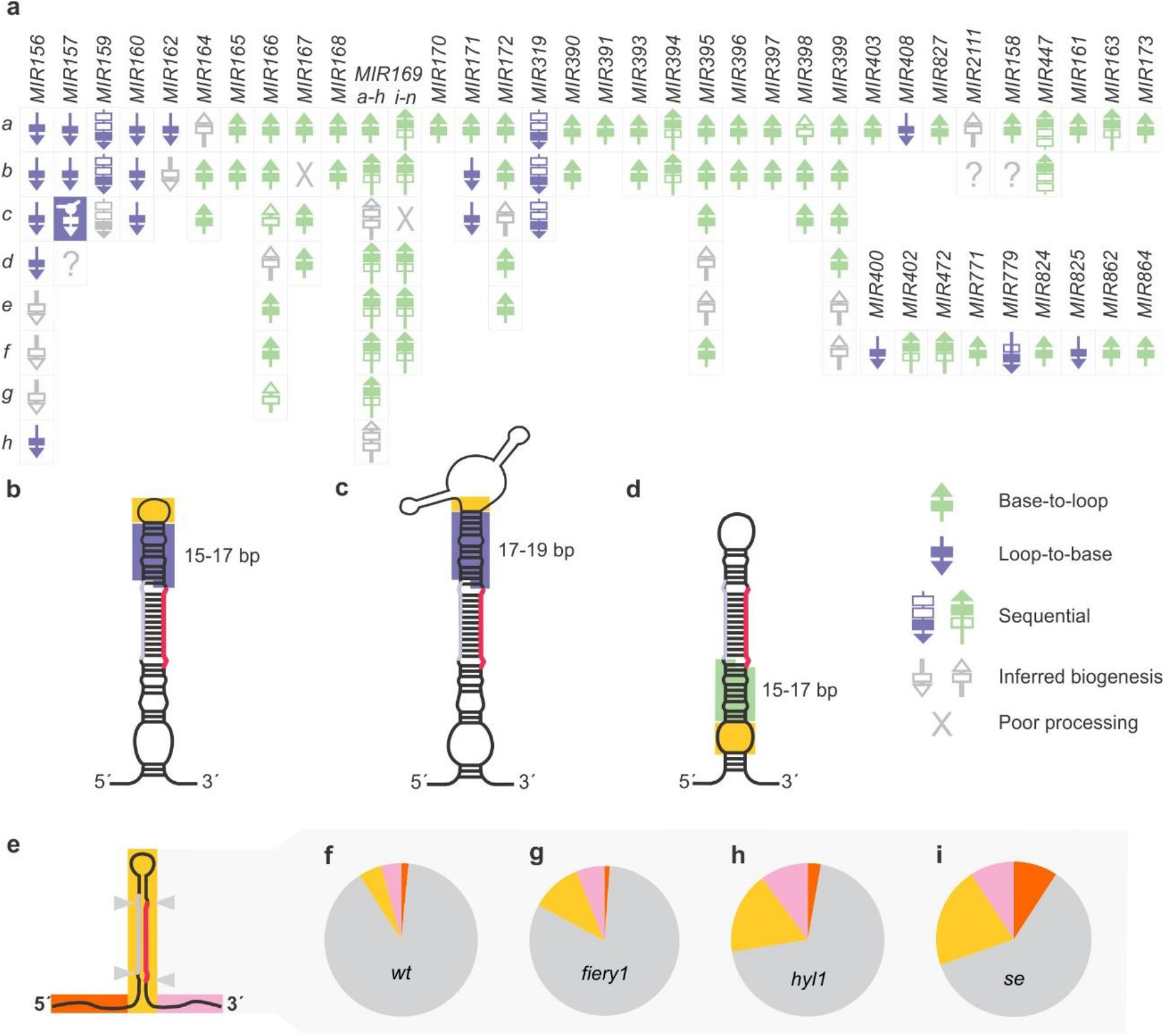
*MIRNA* processing precision in Arabidopsis plants. **(a)** Schematic summarizing the processing of Arabidopsis *MIRNAs.* **(b)** Schematic representation of a precursor processed from the loop, which has a terminal loop (yellow box) followed by a 15-17 bp stem of dsRNA (purple box). **(c)** Schematic representation of *MIR157c,* which has a terminal branched loop (yellow box) followed by a 17-19 pb dsRNA stem above the miRNA (purple box). **(d)** Schematic representation of a precursor processed from the base, which has an internal loop (yellow box) followed by a stem 15-17 bp dsRNA (green box). **(e)** Localization of cleavage sites on Arabidopsis primary *MIRNAs.* Grey triangles represent cuts flanking the miRNA/miRNA*. **(f-i)** Distribution of cuts in primary transcripts in wild-type plants **(f)** and *fiery1* **(g)**, *hyl1* **(h)** and *se* **(i)** mutants. Sector colors correspond to **(e)**, the gray sector corresponds to cuts flanking the miRNA/miRNA*.

We classified the cuts in the *MIRNA* precursors depending on whether they were flanking the miRNA/miRNA*, other regions of the foldback, and upstream or downstream the stem loop (Figure 7e). Even though miRNA biogenesis can proceed through different modes, we observed that in nearly all cases, cuts were flanking the miRNA/miRNA* duplexes in wild-type samples (Figure 7f, and Appendix), suggesting that the miRNA processing machinery is in general accurate. An exception was *MIR169* family members, which are processed sequentially in a base-to-loop direction. Some of these *MIRNAs* were shown to be transcribed in tandem with other members of the same family (44). We noticed that *MIR169k,* a precursor that is downstream of a tandem with *MIR169j* had poor processing precision (Supplementary Figure 5). In contrast, we observed an accurate processing of the precursors upstream of the tandem.

### Unproductive cuts detected in *MIRNA* precursors in Arabidopsis mutants

Analysis of *fiery1* data revealed a minor increase in the frequency of erroneous cuts (Figure 7g) with respect to wild type (Figure 7f). We explored in more detail the erroneous cuts in the stem-loop precursor. We found additional cuts that were likely triggered by terminal branched loops (see *MIR393a, MIR396a, MIR403* and *MIR864* in Appendix), in agreement with previous results showing branched loops guiding unproductive cuts that destroy *MIRNA* precursors (29). Interestingly, we identified additional cleavage sites in *fiery1,* which are compatible with the recognition of additional alternative structural determinants (Figure 8). In *MIR165a, MIR170, MIR395c,* and *MIR157b* we found a cleavage site 15-16 bp distal to an internal bubble in the region below the miRNA/miRNA* (Figure 8a-d).

**Figure 8.**
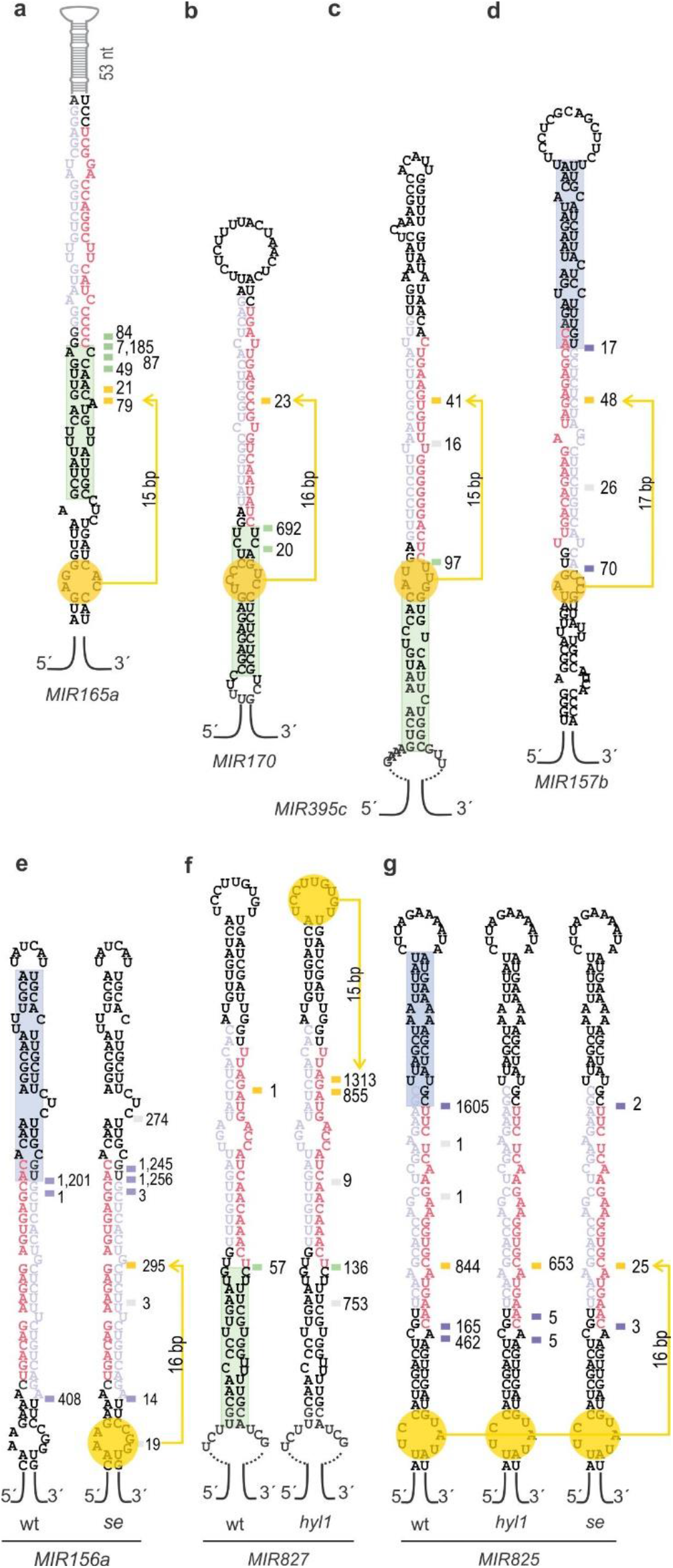
Unproductive processing of *MIRNAs* detected in Arabidopsis mutants. **(a-g)** Predicted secondary structure of **(a)** *MIR165a,* **(b)** *MIR170,* **(c)** *MIR395c,* **(d)** *MIR157b,* **(e)** *MIR156a,* **(f)** *MIR827,* and **(g)** *MIR825.* The structural determinant recognized during processing in wild-type plants is indicated with a green (base-to-loop) or purple (loop-to-base) rectangle. Alternative structural determinants are indicated in yellow. **(a-d)** Cuts indicated correspond to *fiery1* samples.

Next, we analyzed *hyl1* and *se* mutants that have key roles in miRNA processing efficiency (4,5,8,9,12). To do this, we also prepared SPARE libraries from these mutants. Even though *hyl1* and *se* demonstrate a strong reduction in miRNA biogenesis, we were able to detect processing intermediates in these mutants. However, we also noticed an important increase in the frequency of erroneous cuts (Figure 7h-i). Analysis of localization of the cuts along the miRNA primary transcripts revealed an increase of cuts in the 5’ end and 3’ end region of the precursors, with respect to wild-type samples (Figure 7e-i). Furthermore, we noticed that in certain precursors there were additional cuts present in *hyl1* and/or *se* that were also consistent with the recognition of additional structural determinants (Figure 8). A cleavage site in *MIR156a* was consistent with the recognition of an internal bubble below miR156/miR156* in *se* (Figure 8e), while we observed a cut in *MIR827* ~16 bp away from the small terminal loop in *hyl1* mutants (Figure 8f). Neither wild-type nor *fiery1* samples showed these cuts (Figure 8ef, Appendix). In addition, we observed an enhanced cut in *MIR825* in both *hyl1* and *se* mutants with respect to wild-type plants (Figure 8g). The results suggest that alternative or cryptic structural determinants can be active during miRNA biogenesis in certain conditions or genetic backgrounds.

## Discussion

The position of the first cleavage site in *MIRNA* precursors is essential to determine the sequence of the mature miRNA. While the second cut uses the staggered ends of a processed precursor committed in the miRNA biogenesis pathway, the first cleavage must rely on structural cues that guide the processing complex to the correct position. A common aspect between the animal and plant processing machinery is that the processing complex defines the initial cut in a primary miRNA transcript at a distance from a ssRNA-dsRNA junction and into a stem region. However, plant *MIRNA* precursors are more variable than their animal counterparts and are, in turn, processed by different modes. In certain cases, the processing machinery recognizes an internal bubble and cuts 15-17 bp away in a lower stem to release a fold-back precursor (22–25,29,33). In other cases, a small terminal loop and a 15-17 bp upper stem above the miRNA/miRNA* are recognized in a loop-to-base processing mode [(21), this work]. In both cases, the processing complex acts as a molecular ruler recognizing a ssRNA-dsRNA transition, between a stem and an internal bubble or a terminal loop.

It has been shown that primary miRNA transcripts with branched loops can also be recognized and cut unproductively by DCL1 in plants (29). This recognition occurs on top of the biogenesis of the miRNA as pathway that degrades the *MIRNA* precursor (29). Here, we report that this pathway can lead to the productive biogenesis of mature miRNAs in wild-type plants, as members of the *MIR157* family are processed via the recognition of branched loops. Based on these results, terminal branched loops can be added to the toolkit of structural determinants recognized by DCL1 complexes, highlighting the plasticity of the small RNA pathways in plants. Whether the processing mode of *MIR157c* has a specific biological role remains to be elucidated. Since the efficiency of miR157c biogenesis was improved after replacing the terminal branched loop by a small loop, the processing efficiency might also vary with the different biogenesis modes.

Interestingly, we detected some differences in the processing modes of *MIRNA* precursors. The first cut by DCL1 is more flexible in those precursors processed from the loop than in those initiating from the base. This difference in turn causes variation at the level of the small RNAs. It has been shown that variation in mature miRNA sequences of miR168 causes the sorting of the miRNA into different AGOs and modify its activity (45, 46). In this case, heterogeneity in miR168 sequences is the result from the flexibility of the miRNA/miRNA* region, which affects the position of the second cut by DCL1 (45). Our results show that isomiRs can also be generated by variability in the position of the first cut, especially in those precursors processed from the loop. Actually, isomiRs deriving from *MIR157c* are quite frequent, representing more than 30% of the mature sequences, so its plausible that the structural flexibility conferred by the terminal branched loop contributes to this variation.

Both HYL1 and SE participate in the different biogenesis modes of plant miRNAs, as mutations in their genes result in a global decrease of miRNA levels (4,5,8,9). These proteins have been implicated in several processes including processing precision (12,14,16,47,48), strand selection (49) and splicing (10, 50). *In vitro* processing studies using *MIR167b* and *MIR166g* precursors have shown that DCL1 generates more incorrect cuts in the absence of HYL1, especially near the end of the transcript (12, 14). However, *in vivo* studies of wt and *hyl1* small RNA libraries revealed that incorrect small RNAs generated from other regions of the precursors represent less than two percent of the total population (47, 48), suggesting that additional factors play a role during miRNA processing *in vivo.*

The complex landscape of precursor structures and biogenesis modes poses a major challenge to the processing machinery. A single miRNA primary transcript harboring an imperfect fold-back structure may contain several regions that can potentially initiate miRNA processing, such as an internal bubble followed by dsRNA segments and a small or branched terminal loop. Yet, nearly all the cuts found in the precursors of wild-type plants were flanking the miRNA/miRNA*, indicating a high degree of specificity. The analysis of processing intermediates in *fiery1, hyl1* and *se* mutants showed misplaced cuts in the precursors at sites that are coincidental with the recognition of cryptic structural determinants. In the case of *fiery,* that has a minor increase in additional cuts, detection of misplaced cuts could be related to the absence of XRNs that rapidly degrade misprocessed precursors in wild-type plants. On the other hand, HYL1 and SE might have additional functions aiding DCL1 in finding the correct structural determinants in the precursors. In most cases, the cuts that we detected in *MIRNA* precursors in *hyl1* and *se* were 15-17 bp from the ssRNA/dsRNA junctions, indicating that the processing complex can still function as a molecular ruler in the absence of HYL1 or SE. Recent findings in animals showed that the RNase type III DROSHA acts as a molecular ruler to determine the position of the first cut during animal miRNA biogenesis, 11 bp away of the ssRNA/dsRNA transition in the precursor stem (51). Therefore, it is possible that the same ability resides in plant DCL1 proteins, with HYL1 and SE having a major impact on miRNA biogenesis efficiency and additional roles helping DCL1 to identify the correct structural determinants.

## Supplementary Material

### Appendices

**Appendix**. SPARE reads mapped to Arabidopsis *MIRNAs* for wild-type, and *fiery1* plants.

### Supplementary Figures

**Supplementary Figure 1**. Original images of miR157 small RNA gel blot.

**Supplementary Figure 2**. Schematic illustrating the processing intermediates detected by the SPARE method.

**Supplementary Figure 3**. Alternative RNA folding structures of *MIR157c.*

**Supplementary Figure 4**. IsomiRs detected in wild-type plants due to variation in the first cleavage site.

**Supplementary Figure 5**. Processing intermediates of tandem *MIR169i-n* family members.

### Supplementary Tables

**Supplementary Table 1**. SPARE primers and mix for cDNA synthesis.

**Supplementary Table 2**. Expressed sequences from binary vectors.

**Supplementary Table 3**. Normalized SPARE reads for wild-type and *fiery1* libraries.

**Supplementary Table 4**. Small RNA sequences used to calculate miRNA accuracy.

## Author contributions

B. M. and J.F.P. designed the experiments, B.M. performed the miRNA experiments, constructed the RNA library’s aided by S.A. and analyzed the data, U.C. processed the raw data from the SPARE library. I.P.S did the structural mapping for *MIR171b* and aided B.M. with the structural mapping of *MIR157c.* B.M., R.M.R., B.C.M., C.H., and J.F.P. analyzed the data. B.M. and J.F.P. wrote the article.

## Acknowledgments

We thank Andrea Gamarnik for the V1 enzyme, and members of the J.F.P lab for comments and discussions.

